# Multi-task deep learning for concurrent prediction of protein structural properties

**DOI:** 10.1101/2021.02.04.429840

**Authors:** Buzhong Zhang, Jinyan Li, Lijun Quan, Qiang Lyu

## Abstract

Protein structural properties are diverse and have the characteristics of spatial hierarchy, such as secondary structures, solvent accessibility and backbone angles. Protein tertiary structures are formed in close association with these features. Separate prediction of these structural properties has been improved with the increasing number of samples of protein structures and with advances in machine learning techniques, but concurrent prediction of these tightly related structural features is more useful to understand the overall protein structure and functions. We introduce a multi-task deep learning method for concurrent prediction of protein secondary structures, solvent accessibility and backbone angles (*ϕ, ψ*). The new method has main two deep network modules: the first one is designed as a DenseNet architecture a using bidirectional simplified GRU (GRU2) network, and the second module is designed as an updated Google Inception network. The new method is named CRRNN2.

CRRNN2 is trained on 14,100 protein sequences and its prediction performance is evaluated by testing on public benchmark datasets: CB513, CASP10, CASP11, CASP12 and TS1199. Compared with state-of-the-art methods, CRRNN2 achieves similar, or better performance on the prediction of 3- and 8-state secondary structures, solvent accessibility and backbone angles (*ϕ, ψ*). Online CRRNN2 applications, datasets and standalone software are available at http://qianglab.scst.suda.edu.cn/crrnn2/.

## 1. Introduction

Proteins are composed of different proportions of amino acids (residues) assembled using peptide bonds into unique 3-D structures. The protein’s functions are largely determined by its tertiary structures. Understanding protein structural properties is critical for the study of protein 3D structure and functions. In proteins, the secondary structure is the local conformation of a protein’s polypeptide backbone, which is formed by the patterns of hydrogen bonds between backbone amide and carboxyl groups. Hydrogen bonding is correlated with other structural features, such as dihedral angles. The conformation of a protein’s backbone structure can be described continuously by the two torsion angles *ϕ* and *ψ*.

The solvent-Accessible Surface Area (ASA) is also an important structural features that can measure the degree of a protein’s exposure in its folded state. The specific value is determined by the position of the side-chain group. Solvent accessibility is closely involved with structural domains identification [1],fold recognition [2], binding region identification [3], protein-protein interactions [4], and protein-ligand interactions [5].

The Anfinsen’s experiment [6] has been shown that the structural properties of a protein are encoded in its primary sequence. As a practical application of it, predicting the three-dimensional structure of a protein given only its primary sequence is still a challenging problem with unsatisfactory solution performance. As an alternative approach, predicting intermediate structural properties can give more insight into the native structures. In recent decades, many computational methods have been developed to predict protein structural features from sequences, including secondary structures, backbone angles, disorder, solvent accessible surface area, contact maps [7, 8, 9, 10, 11].

The best-known protein structure prediction subproblem[12] is the prediction of protein secondary structure, where the structures are characterized into 3-states [13]or 8-states [14]. Three-state prediction from protein sequences has been intensively studied using many machine learning methods, including support vector machines [15, 16], hidden Markov models [17, 18], artificial neural networks [19, 20, 21, 22],and the probability graph models [23, 24]. Recently, research has shifted to focus on the complex problem of 8-state structure prediction for a more complex model trained by more sequences can provide better generalization. Many achievements have been accomplished by deep learning techniques, such as the SC-GSN network [25], bidirectional long short-term memory (BLSTM) [26, 11, 27], deep conditional neural fields [28], DCRNN [29], conditioned Convolutional Neural Network(CNN)[30], Deep inception-inside-inception (Deep3I) networks [31], CRRNN[32], NetSurfP-2.0 [27], SPOT-1D [33].

The methods for ASA prediction are from discrete classification [34, 35], relative continuous value [10, 36] to real value of surface area [11, 33]. Other methods have been developed to predict *ϕ* and *ψ* as both discrete [37] and continuous values [8, 38, 39, 11].

The protein’s tertiary structure is coherently related with its secondary structure, backbone angles and ASA. Jointly predicting structural features is an effective way to improve model generalization. In 2005, Wood and Hirst[8] predicted protein secondary structure and *ψ* dihedral angle. Cheung *et al*. [40] employed Bayesian inference to predict the backbone dihedral angles *ϕ, ψ* and secondary structure. Faraggi et al [22] developed a multi-step neural network for predicting secondary structure in an iterative manner given predicted solvent accessibility and backbone torsion angles. Heffernan *et al*.[41, 11] used a Bidirectional Recurrent Neural Network(BRNN) to predict 3-state secondary structure, ASA, backbone torsion angles and contact numbers in a similar manner.

Recently, many deep learning methods have been successfully applied to predict protein structural properties. These methods, such as NetSurfP-2.0 and SPOT-1D are more generalizable for being trained by more than 10k sequences, using different types of neural networks such as BRNN, CNN and ResNet, with input features including PSSM, HMM [42]. However, the model scale of NetSurfP-2.0 and SPOT-1D is very large, and their very large of parameters need more computing resources.

This paper proposed a new method called CRRNN2 for concurrently predicting the one dimensional protein structural properties of secondary structure, ASA, and backbone angles by deep multi-task learning. The CRRNN2 architecture is composed of DenseNet-based[43] BRNNs, one dimensional CNNs and a variant of the Inception network[44]. CRRNN2 is trained by 14600 sequences and tested by public test datasets such as CB513, CASP10, CASP11, CASP12 and TS1199. Compared with existing models, CRRNN2 is more lightweight but the generalization is comparable or higher.

## 2. Method

### 2.1 CRRNN2 Architecture

Protein structures are determined not only by local residues, but also are strongly affected by long-range residues. As described above, BRNN or CNN-based models or hybrid networks effectively capture protein long-range and local structural features. The CRRNN2 model is consisted of DenseNet[43] based BRNNs, one-Dimensional CNNs [32] and a variant of Google Inception[44], as shown in Fig.1. A one-Dimensional CNN with kernel size one (denoted as 1D1 CNN), 85 filters is used to extract the input features from PSSM, sequence coding and HMM profile respectively. After a concatenation operation(Concat), the 255-dimensional features are fed to DenseNet based BRNNs(DenseNet-BRNN) and variant of Inception(V-Inception) respectively. The outputs of DenseNet-BRNN and V-Inception are concatenated again and then fed to two fully connected(FC) layers. The predicted values of the model output are Q8, Q3, Q3 2, RSA, *sin*(*ψ*), *cos*(*ψ*), *sin*(*ϕ*) and *cos*(*ϕ*).

**Fig. 1.**
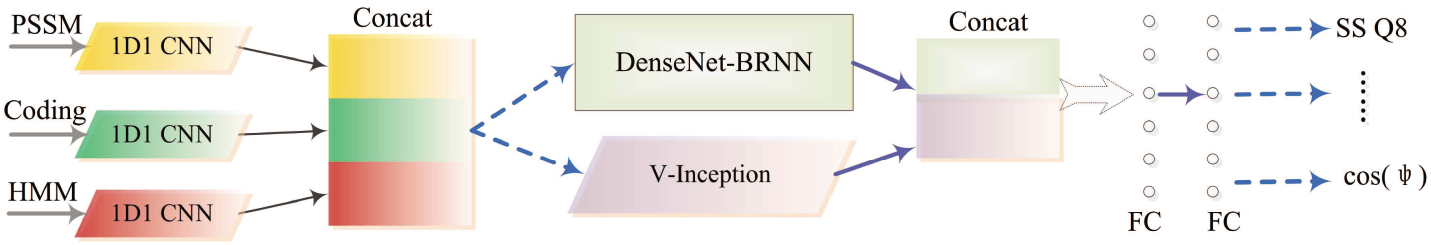
The model architecture of CRRNN2.

The V-Inception network is more detailed shown in Fig.2. The one-dimensional CNN is replaced the two-Dimensional CNN. The kernel size are one(1D1), three(1D3) and five(1D5), and 85 filters are used respectively. The max pooling operation is replaced by directly connecting the input data. The operations indicated by dotted lines are concatenating, dimension reduction and dropout(*p*=0.5). Formalized descriptions are presented in Equation(1). The 1D1 CNN with 150 filters is used to reduce the number of dimensions. A weight constraint of Dropout (*p*=0.5) is used to avoid overfitting. These three operating procedures together are named as converted layer.

**Fig. 2.**
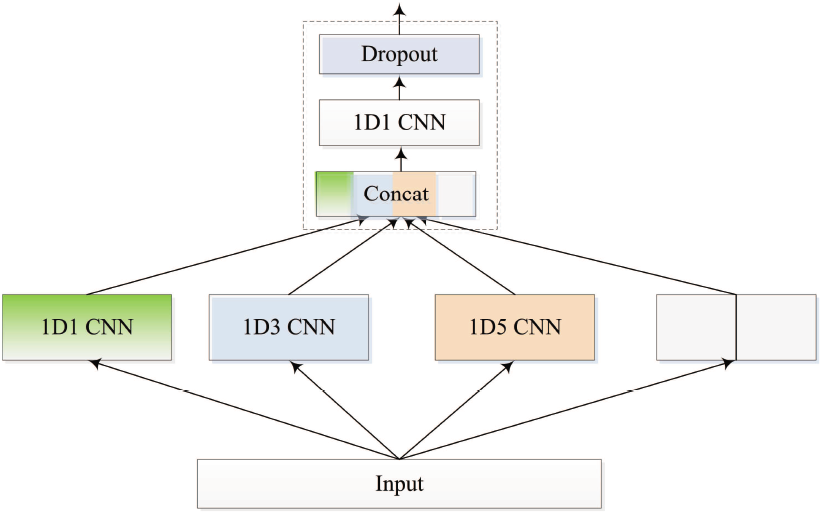
Variant of Inception network. one-dimensional CNNs are used and the activation function is ReLU. The kernel size is one(1D1), three(1D3) and five(1D5), and 85 filters are used respectively. Max pooling operation is replaced by directly connecting the input data. 1D1 CNN with 150 filters is used for reducing the number of dimensions.

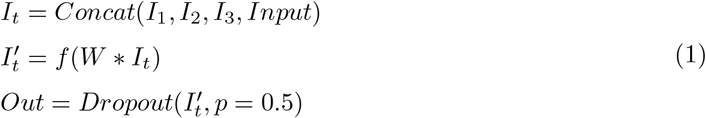

The DenseNet-based BRNNs are illustrated in Fig.3. Three bi-directional recurrent neural network layers are stacked. Like CRRNN, the merging computation in the first BRNN layer is concatenated, and the others are summed when the forward-computed result *F*_*t*_ is merged with the backward result *B*_*t*_. Each output from a BRNN layer is connected with the converted layer. In the first, second and third BRNN layer, 250,500 and 500 units respectively are used in the unidirectional recurrent neural network. The dimensionality of every layer output is 500 and the number of CNN filters in the converted layer is also 500. The output activation function of each BRNN layer is tanh function.

**Fig. 3.**
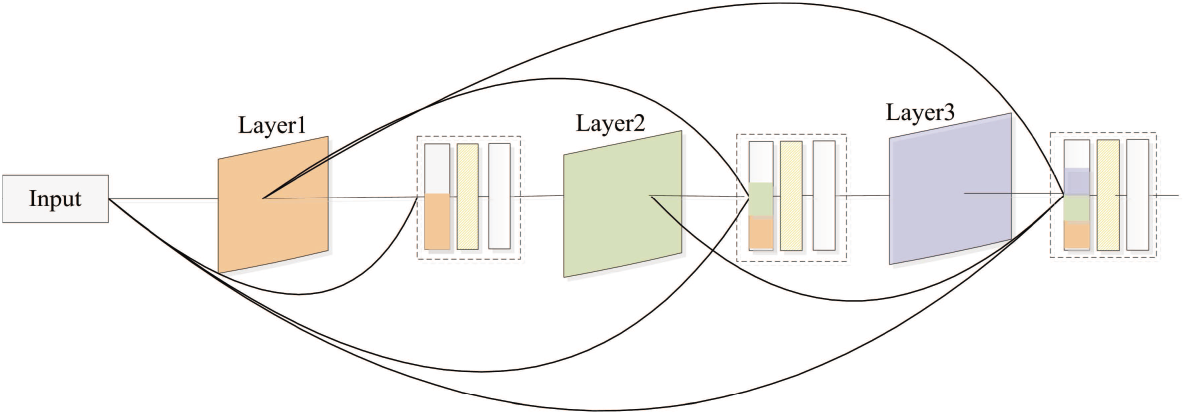
DenseNet-based BRNN.

Zhang *et al*’s description and experiment[45] has been proved that neural network performance is slightly affected when the previous state *h*_*t*−1_ in the input gate and output gate is removed from Long short-term memory network. A simplified GRU network(GRU2) is used to act as an unidirectional recurrent neural network in CRRNN2 and is described by Equation(2):

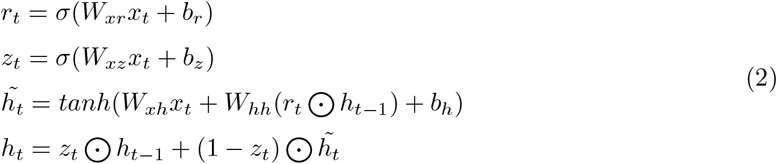

where *σ* is the sigmoid function, ⊙ represents an element-wise multiplier. 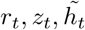 and *h*_*t*_ are the reset gate, update gate, internal memory cell activation vectors and output, respectively.

### 2.2 Multi-task learning

By sharing representations among related tasks, multi-task learning can enable model to generalize better than the original single task [46]. As a protein’s structure is closely related with its secondary structure, backbone angles and ASA, concurrently predicting these features is helpful to obtain a better representation from one model. The hard parameter sharing strategy [46] is employed in CRRNN2. Each task shares the input and hidden layers except for the output layer.

The output of CRRNN2 consists of the predicted secondary structure labels 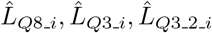 the RSA values the 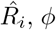, angle’s sine and cosine values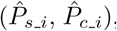, and the *ψ* angle’s sine and cosine values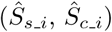. The joint loss function is formulated as follows:

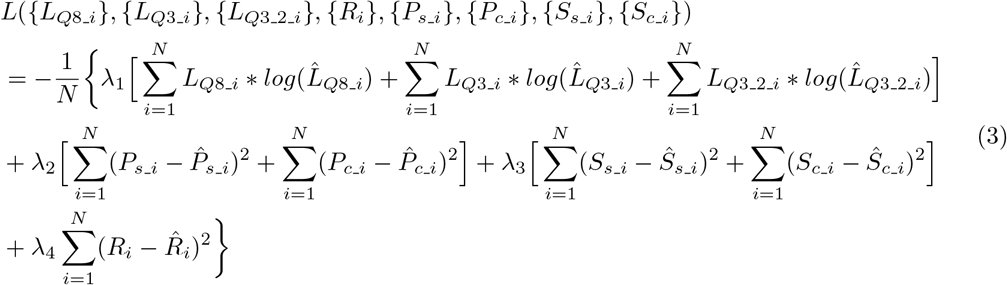

In Equation(3), *λ*_1_ = 1, *λ*_2_ = *λ*_3_ = 1.4 and *λ*_4_ = 4.0 which is obtained by grid search. Different loss functions are used for different learning tasks. The softmax cross-entropy loss function is used for multi-classification task learning of Q8, Q3 and Q3 2 predictions. The Mean square error (MSE) function is used for trigonometric function fitting and RSA real value fitting.

## 3. Results and discussion

### 3.1 Datasets

The training dataset contains 14458 polypeptide chains obtaining from four CullPDB datasets[47] and cons-PPISP’s[48] training dataset. The degree of identity of the CullPDB datasets is no greater than 25% (first three datasets) or 30% (the final dataset) similarity, the resolution better than 3.0 Å and an R-factor is 1.0. As in the previous work[32], the sequence which length is more than 700 or less than 50 is removed.

Sequences with a degree of similarity greater than 40% with those in testing datasets were removed from the candidate training dataset by the Cd-hit [49] software, and similar proteins from different data sources were also removed from the training sequences. Finally, 358 sequences are selected as a validation dataset (VD358) for evaluating the model or for early stopping during training. This left 14,100 sequences (TR14100) to be used for training.

Four public test datasets (CASP10 (101 sequences), CASP11, CASP12, and TS1199) are used to evaluate the performance of backbone angles and solvent-accessible area prediction. Five test datasets (CB513, CASP10 (123 sequences), CASP11, CASP12, and TS1199) that were used in previous work [32] are also used to evaluate the model generalization capability for secondary structure (Q3 and Q8) prediction.

### 3.2 Input vectors and outputs for multi-task learning

Three types of features are selected, each consisting of a protein sequence coding[32] and two evolutionary profiles from three iterations of PSI-BLAST [50] with a default E-value and HHBlits [42] with default parameters. Each residue in the sequence is characterized by 22-dimensional sequence coding, 20-dimensional Position-specific scoring matrix(PSSM), and 30-dimensional HMM profile.

For each sequence in this study the real values of solvent accessible area, 3-and 8-state secondary structure, and the backbone dihedral angles (*ψ, ϕ*) are calculated by DSSP[14] software. The ASA of each residue is normalized to [0, 1] by *ASA*_*i*_*/Max*_*ASA*_. The backbone dihedral angles *ψ* and *ϕ* are represented by a couple of trigonometric functions *sin*(*α*) and *cos*(*α*). The predicted angle is recovered at the network output by 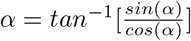. For comparison with existing methods, two mapping patterns of 8-state labels condensed to 3-state are used. The first pattern named Q3 is H(8 state H), E(8 state E) and C (everything else). Another pattern named Q3 2 is H(8 state G, H and I), E(8 state B and E), and C(others).

### 3.3 Training setup

In our experiments, the size of the fist full connected layer is 1024. The second full connected layer is each task learning layer. An Adam optimizing function is used for training the model with the default setting parameters. The default learning rate is initially set at 0.0004 with a decreasing ratio 0.2, whereas the loss on the validation dataset VD315 does not decrease after more than 20 epochs.

CRRNN2 model is implemented in Keras, which is a publicly available deep-learning software. The weights in CRRNN2 are initialized using default values, and CRRNN2 is trained on a single NVIDIA RTX 2080 Ti GPU with 11GB memory.

### 3.4 Impact of different input features and model architectures

The performance of the CRRNN2 model using different RNN units and different groups of feature profiles for the 1D structural properties prediction were rigorously examined using the training and validation dataset presented in this work.

The impact of different groups of input features is firstly analyzed. Q8 prediction performance is compared using different combinations of input features on validation dataset. the iterating procedure is illustrated in Fig.4. PSSM profile is assumed to be the primary feature for many previous studies have verified it. The predicting accuracy is about 74%. When the sequence coding or HMM profile is included, the accuracies improve to 74.6% and 77.2% respectively. When all three kinds of features are integrated, the model achieves its highest score (about 78%).

**Fig. 4.**
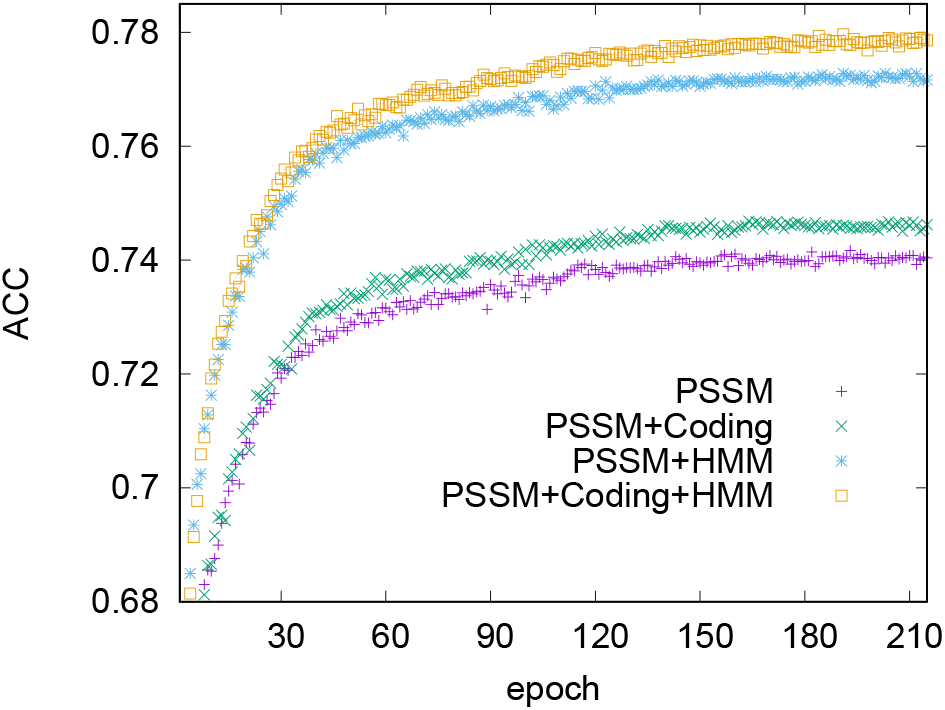
The Q8 prediction performance comparison of the iterative procedure using different input features on the validation dataset.

The impact of different RNN units in the CRRNN2 model is also analyzed. The performance of the CRRNN2 model using GRU, LSTM, SLSTM and GRU2 unit is compared. The prediction results for the four deep architectures on the VD315 dataset are reported in Table 1. As shown in Table 1, the performance of CRRNN2 using GRU, LSTM, SLSTM[45] and GRU2 is not obviously different. To evaluate the model generalization further, the loss variation of the various models is compared and is illustrated in Fig. 5. The model using SLSTM and GRU2 obtained smaller less on the validation dataset. When the least loss point was reached, the loss of models using LSTM and GRU then increased speedily, but the those of the models using SLSTM and GRU2 increased more slowly. In contrast, the model using SLSTM or GRU2 is more generalizable. The numbers of parameters in the CRRNN2 models using GRU, LSTM, SLSTM, and GRU2 units are 9.62 million, 11.87 million, 8.5 million and 7.37 million respectively. The number of parameters in the CRRNN2 using GRU2 was the smallest, making the model more lightweight.

**Table 1.** Performance comparison of multi-structural properties prediction on the VD358 dataset.

| Method | Parameters(M) | Q8 | Q3 | Q3.2 | ASA(PCC) | $\phi$(MAE) | $\psi$(MAE) |
| --- | --- | --- | --- | --- | --- | --- | --- |
| CRRNN2_GRU | 9.62 | 0.775 | 0.899 | 0.88 | 0.801 | 18.82 | 24.5 |
| CRRNN2_LSTM | 11.87 | 0.776 | 0.899 | 0.879 | 0.801 | 18.81 | 24.3 |
| CRRNN2_SLSTM | 8.5 | 0.778 | 0.901 | 0.882 | 0.802 | 18.79 | 24.12 |
| CRRNN2_GRU2 | 7.37 | 0.777 | 0.902 | 0.882 | 0.803 | 18.8 | 24.23 |

**Fig. 5.**
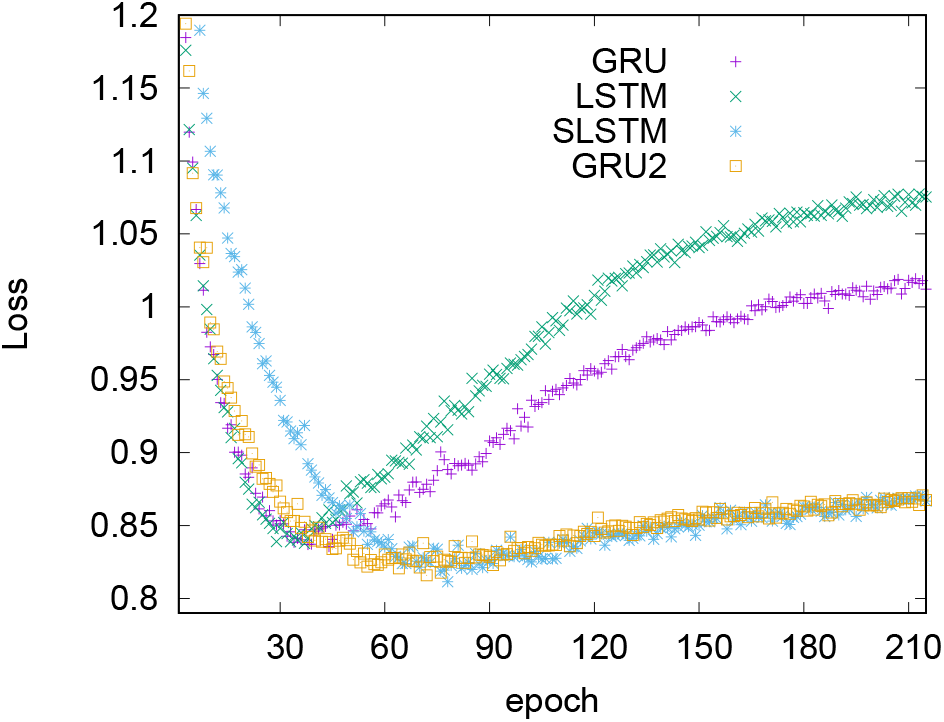
The variation in losses between CRRNN2 models using bidirectional GRU, LSTM, SLSTM and GRU2 on the VD358 validation dataset.

The last evaluation was intended to investigate whether the new deep architecture network (CRRNN2) outperformed the previous network (CRRNN) for the secondary structure prediction.

To achieve a fair comparison, CRRNN was trained by TR14100 with the same input features as CRRNN2. The predicted results are reported in Table 2 and 3 and are denoted as CRRNN^*a*^. CRRNN^*a*^ achieved 72.9%, 74.5%, 73.4%, and 71.7% of Q8 prediction and 87.6%, 87.4%, 86.7%, and 84.8% for Q3 prediction on CB513, CASP10, CASP11 and CASP12 respectively. Compared with the original CRRNN model, although the performance of CRRNN^*a*^ is improved, it’s still worse than that of CRRNN2. CRRNN2 has fewer parameters but provides better generalization. The comparison showed that model generalization performance is strongly affected by the deep architecture.

**Table 2.** Performance comparison on Q8 prediction between CRRNN2 and existing methods (%)

| method | CB513 | CASP10 | CASP11 | CASP12 |
| --- | --- | --- | --- | --- |
| DeepCNF | 68.3 | 71.8 | 71.7 | 69.4 |
| DCRNN | 70.4 | 73.9 | 71.2 | 68.8 |
| MUFOLD-SS | 70.5 | 76.5 | **74.5** | 72.1 |
| NetSurfP-2.0 | 72.3 | - | - | 71.1 |
| NetSurfP-2.0* | 72.5 | 74.4 | 72.6 | 71.1 |
| CRRNN | 71 | 74.3 | 72.3 | 70.1 |
| CRRNN<sup>a</sup> | 72.9 | 74.5 | 73.4 | 71.7 |
| CRRNN2 | **74.9** | 75.9(**77.5**) | 74.3 | **72.9** |

\*Results are obtained by our experiments. NetSurfP-2.0\* is our re-implemented model. Boldface numbers indicate best performance.

**Table 3.** Performance comparison on Q3 prediction between CRRNN2 and existing methods (%)

| Method | CASP10 | CASP11 | CASP12 | CB513 |
| --- | --- | --- | --- | --- |
| PSIPRED | 81.2 | 80.7 | 80.5 | 79.2 |
| JPRED | 81.6 | 80.4 | 78.8 | 81.7 |
| DeepCNF | 84.4 | 84.7 | 83.2 | 82.3 |
| MUFOLD-SS | 86.5 | 85.2 | 83.4 | 82.7 |
| NetSurfP-2.0 | - | - | 82.4 | 85.4 |
| NetSurfP-2.0* | 87.3 | 85.2 | 84.4 | 87.1 |
| CRRNN | 86.7 | 84.7 | 83.8 | 85.7 |
| CRRNN<sup>a</sup> | 87.4 | **86.7** | 84.8 | 87.6 |
| CRRNN2 | **88.3** | 86.5 | **85.5** | **88.4** |

\*Results are obtained by our re-implemented experiments.

### 3.5 Comparison on independent test datasets

To evaluate the model generalization of CRRNN2, its performance on the secondary structure prediction is compared with that of other state-of-the-art methods: the DeepCNF [28], DCRNN [29], MUFOLD-SS [31], CRRNN^*a*^ and NetSurfP-2.0 [27]. Table 2 shows a comparison of the results for Q8 prediction accuracy. The proposed model achieved 74.9%, 75.9%, 74.3%, 72.9% accuracy on CB513, CASP10, CASP11 and CASP12 test dataset respectively. The NetSurfP-2.0^∗^ method is re-implemented in our experiment and trained using TR14100 on four Nvidia GTX 1080ti GPUs. About 34.99 million parameters are existed in the re-implemented model. The CASP10 dataset used in MUFOLD-SS method contains 101 sequences. The prediction accuracy of our method on this dataset is 77.5% which is indicated in the bracket, Table 2. From Table 2, it can be concluded that the present CRRNN2 model obtains more generalizable performance.

Q3 prediction performance is also compared and shown in Table 3. The CRRNN2 model obtained 88.3%, 88%, 86.5%, and 88.4% accuracy on CASP10, CASP11, CASP12 and CB513 test datasets respectively. On another mapping mode of Q8 condensed to Q3(Q3 2), the CRRNN2 model obtained 87.3%, 86%, 84.1%, 88.1% and 87.1% accuracy on CASP10, CASP11, CASP12, CB513 and TS1199 test set respectively. Fig.6 compares the accuracy of Q3_2 secondary structure prediction at individual amino acid levels with CRRNN^*a*^, SPIDER3, and SPIDER2 on the TS1199 test dataset, indicating highest accuracies (86.5%). As the theoretical limit(88-90%) of three-state prediction[12], the performance of CRRNN2 has been reached or approximated the theoretical limit on the first mapping mode.

**Fig. 6.**
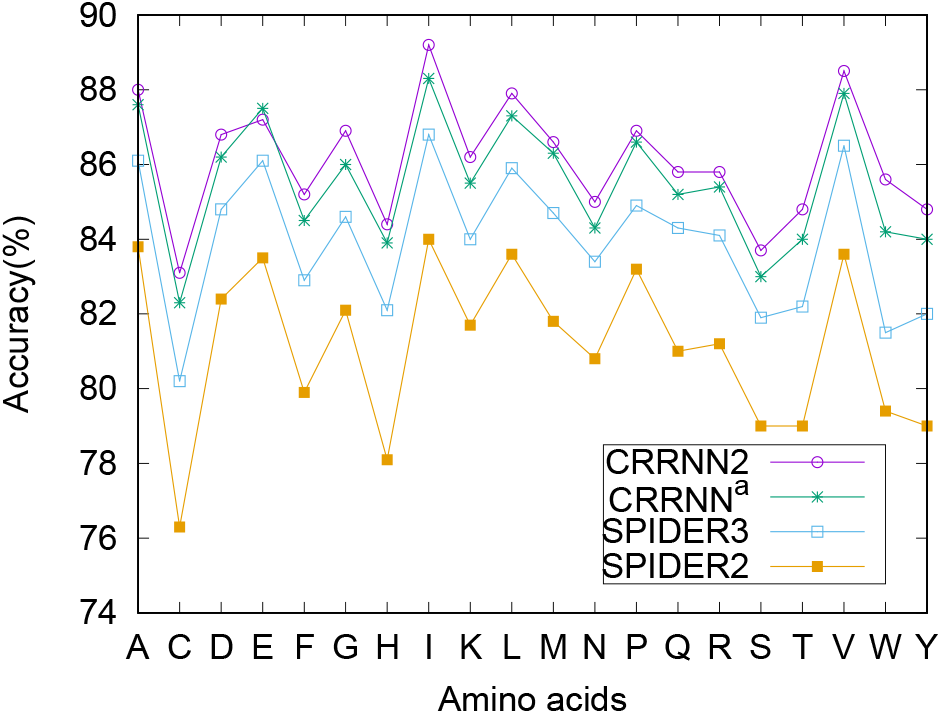
The accuracy of Q3 2 secondary structure prediction for individual amino acids as compared with SPIDER2, SPIDER3,CRRNN^*a*^ and CRRNN2, on the TS1199 dataset.

The backbone angle prediction was evaluated by the mean absolute error (MAE) and Pearson correlation coefficient(PCC) between the truth and the predicted angles for each residue. Considering the periodicity in angles, the MAE for prediction of residue *i* is defined as the smaller value of *d*_*i*_ and 360^°^ *− d*_*i*_, where *d*_*i*_ is the average absolute difference between predicted angles (P) and experimentally determined angles (E), and P has been transformed into [*−*180^°^, 180^°^] [38].

Ninety-nine sequences out of 123 in CASP10, 85 out of 85 in CASP11, and 40 out of 40 in CASP12 are used as public test datasets. Table 4 and 5 compare the results of SPIDER2, SPIDER3, RaptorX-Angle, DeepRIN, and CRRNN2 on the predicting dihedral angle *ψ* and *ϕ*. CRRNN2 outperformed other methods in all cases and also showed better generalization performance. The prediction error of the *ψ* angles is reduced by nearly 2 degrees. However, the prediction error of the *ϕ* angles is just reduced by 1.14-1.47 degree which may have been due to the competitive performance of *ϕ* angle prediction, leaving little room for performance improvement.

**Table 4.** Comparison of MAE(PCC) on angle ψ (PSI).

| Angle | CASP10 | CASP11 | CASP12 |
| --- | --- | --- | --- |
| SPIDER2 | 49.18(0.693) | 48.13(0.699) | 50.59(0.688) |
| SPIDER3 | 31.17(0.824) | 33.05(0.805) | 35.67(0.783) |
| RaptorX-Angle | 34.85(0.8) | 30.14(0.788) | 31.6(0.834) |
| DeepRIN | 25.79(0.865) | 27.96(0.855) | 31.39(0.834) |
| NetSurfP-2.0* | 26.16(0.869) | 28.68(0.854) | 30.88(0.843) |
| CRRNN2 | **24.02(0.878)** | **26.68(0.864)** | **28.9(0.853)** |

NetSurfP-2.0\* is our re-implemented model. Boldface numbers indicate best performance.

**Table 5.** Comparison of MAE(PCC) on angle *ϕ*(PHI).

| Angle | CASP10 | CASP11 | CASP12 |
| --- | --- | --- | --- |
| SPIDER2 | 25.83(0.766) | 26.06(0.756) | 26.85(0.758) |
| SPIDER3 | 20.41(0.831) | 21.38(0.815) | 20.59(0.819) |
| RaptorX-Angle | 21.1(0.818) | 20(0.798) | 20.69(0.788) |
| DeepRIN | 17.39(0.867) | 18.97(0.849) | 20.21(0.828) |
| NetSurfP-2.0* | 18.03(0.862) | 19.63(0.839) | 20.44(0.828) |
| CRRNN2 | **16.56(0.879)** | **18.49(0.851)** | **19.14(0.842)** |

NetSurfP-2.0\* is our re-implemented model. Boldface numbers indicate best performance.

As reported, SPOT-1D is the latest state-of-the-art model. The accuracy on Q8 and Q3 2, the refined segment overlap(SOV)[51] score on Q8 and Q3 2 prediction, and the MAE and PCC on *ϕ* and *ψ* are compared. The ASA(PCC) value of NetSurfP-2.0 in brackets is from the published paper. Table 6 shows a comparison of the results. The comparison shows that the CRRNN2 performance is comparable to SPOT-1D except for predicting ASA values.

**Table 6.** Performance comparison of multi-structural property prediction on CASP12 dataset.

| Method | Q8 | $SOV_{Q8}$ | Q3.2 | $SOV_{Q3}$ | ASA(PCC) | $\phi$(MAE) | $\phi$(PCC) | $\psi$(MAE) | $\psi$(PCC) |
| --- | --- | --- | --- | --- | --- | --- | --- | --- | --- |
| Spider3 | - | - | 0.825 | 0.729 | 0.727 | 20.59 | 0.819 | 31.6 | 0.834 |
| NetSurfP-2.0* | 0.711 | 0.712 | 0.825 | 0.725 | 0.706(0.737) | 20.44(20.0) | 0.828 | 30.88(31.2) | 0.843 |
| SPOT-1D | 0.739 | 0.732 | 0.846 | **0.759** | **0.764** | **18.91** | 0.84 | **27.46** | **0.866** |
| CRRNN2<sup>b</sup> | **0.74** | 0.733 | 0.846 | 0.751 | 0.742 | 19.3 | 0.841 | 28.3 | 0.859 |
| CRRNN2 | 0.729 | 0.735 | 0.836 | 0.724 | 0.747 | 19.14 | **0.842** | 28.9 | 0.853 |

The CRRNN2<sup>b</sup> is trained by the same dataset and input as SPOT-1D. NetSurfP-2.0\* is our re-implemented model and the ASA(PCC) value in brackets is from its published paper.

SPOT-1D is made up of nine groups of models models with different parameters. The one group of models are integrated by an independent classification and regression model respectively. The size of its storage file ranges from 240MB to 1.8GB. The size of SPOT-1D including all files is about 10.8GB, which makes it hard to as standalone software. However, the CRRNN2 is an individual sequence-based model and the file size is only about 28.9MB. And the CRRNN2 model is a solely model which accomplishes the classification and regression tasks concurrently. The parameters of CRRNN2 are fewer and the model is more lightweight.

## 4. Conclusions

The deep learning networks that have been successfully applied in Bioinformatics include deep convolutional neural networks, bidirectional recurrent neural networks, Resnet, DenseNet, and the Inception network. Based on these techniques, an advanced deep learning architecture is developed in this research, providing sequence-based prediction for secondary structure, solvent accessibility, and backbone angles. Three kinds of features from sequence (PSSM, sequence coding and HMM profile) are used for encoding one residue in the sequence and multi-task learning is effective in resolving the problems of protein structural property predictions.

Extensive experimental results have shown that CRRNN2 consistently generated more competitive performance than other state-of-the-art methods. Compared to other heavy-weight models, CRRNN2 is more lightweight and generalizable. Many works such as eCRRNN, Port 5 and SPOT-1D have proven the effectiveness of ensemble learning. CRRNN2 obtained comparable or more generalization performance against these methods. If more CRRNN2 models are assembled, the performance will be come even more effective.

Although multi-tasks learning can predict many structural properties, not all prediction performance can be improved. In a sense, the result represents a compromise among the performance on all prediction tasks. Under the same conditions, the independent performance on dihedral angle prediction is better than that on multi-task learning.

## ACKNOWLEDGMENTS

This research was supported in part by Industry University Research Innovation Fund of Chinese Universities-the new generation information technology innovation project under grant No. 2019ITA01046, Project of provincial Key Laboratory for Computer Information Processing Technology, Soochow University of China under grant No. KJS1934 and the Excellent Youth Scholars Project of Anhui Provincial Universities of China under grant No. gxyq2020029.

